# TFA-defluorinating microbial consortium achieved with analogue enrichment

**DOI:** 10.64898/2026.10.12.758446

**Authors:** Eva Højgaard Jensen, Asal Forouzandeh, Xenia Trier, Charlotte Held Gotfredsen, Kasper Enemark-Rasmussen, Shaban Ahmad, Tue Kjærgaard Nielsen

## Abstract

Trifluoroacetate (TFA) is an ultra-short chain perfluoroalkyl substance (PFAS) and a pollutant of global significance; it is persistent and ubiquitous in the environment across compartments, including food, drinking water, animals, and humans. Concentrations in the environment are rising, concerns over the toxicity of the compound are increasing, and there are currently no efficient ways to remediate TFA pollution, as conventional methods are ineffective at scale under environmental conditions.

Microorganisms possessing dehalogenating enzymes targeting the structurally analogous compound monofluoroacetate (MFA) present a potential solution to degrade TFA microbiologically. Due to previously demonstrated dehalogenation of difluoroacetate (DFA) by dehalogenases, the promiscuity of such enzymes was hypothesized to be extendable to TFA as well. In this study cultures obtained from an environmental sample of PFAS contaminated wastewater sludge were incubated in M9 minimal media supplied with 33 mM acetate and either 33 mM TFA, DFA, or MFA in an iterative manner, followed by testing in a 30-day experiment. Free fluoride (F^-^) was measured at 5 timepoints using a fluoride ion specific electrode (ISE).

One culture designated MFA1C significantly degraded MFA, DFA, and TFA (two-way ANOVA mixed model, p < 0.05). MFA1C released 11.2 ± 0.7 µM F^-^ which constitutes approximately 0.01% of the potential fluoride release from the total 33 mM TFA applied and with the greatest defluorination activity in the first 9 days of incubation. Fluoride release from TFA and formation of DFA and MFA was confirmed with ^19^F-NMR. Genetic sequencing and protein-ligand modelling identified two candidate dehalogenase enzymes from *Achromobacter denitrificans* and *Stenotrophomonas* sp., respectively, with potential capacity to degrade TFA. The findings of this study are a crucial first step towards future potential bioremediation of TFA, and are, to the best of our knowledge, the first to identify a TFA-degrading consortium of microorganisms.

## Introduction

Per- and polyfluoroalkyl substances (PFAS) have been manufactured since the 1930’s and broadly used in industry and consumer products. The unique physical-chemical characteristics endowed by fluorine provide high stability and make them both water- and oil-repellent, which has proven extremely useful in a wide range of applications. However, their persistence also makes them problematic pollutants, earning PFAS the moniker of ‘forever chemicals’ (Berhanu et al., 2023; Buck et al., 2011; DeWitt et al., 2025; Glüge et al., 2020). Although much focus on PFAS has historically been centred on the prevalence and harmful effects of perfluorooctanoic acid (PFOA) and perfluorooctane sulfonic acid (PFOS) (Mueller & Schlosser, 2020), a still persistent but also mobile PFAS has recently garnered increased interest and concern: trifluoroacetate (TFA) (Arp et al., 2024).

TFA is the perfluorinated analogue of acetate. As a C2 perfluoroalkyl acid, TFA is termed an ultrashort-chain PFAS (Arp et al., 2024). With a p*K*_a_ of 0.2-0.5 (Garavagno et al., 2024) trifluoroacetic acid is a strong acid. It will therefore exist in anionic form in the environment as trifluoroacetate. Due to its small size and negative charge TFA is very soluble in water and generally poorly retained in soil (Richey et al., 1997) and is therefore both highly mobile and highly persistent in the environment (Arp et al., 2024; Freeling & Björnsdotter, 2023; Garavagno et al., 2024; Henne et al., 2025). TFA in the environment comes from a variety of point and diffuse sources such as industrial and municipal waste water, agrochemicals, and atmospheric oxidation of refrigerants (Freeling & Björnsdotter, 2023; Scheurer et al., 2017) and is present in a very wide range of environmental compartments. It can be found everywhere from ice cores in the arctic, the oceans, and precipitation, to soil, plant material, food, and beverages, as well as in wildlife and human blood serum (Duan et al., 2020; European Commission, 2017; Freeling et al., 2022; Freeling & Björnsdotter, 2023; Garavagno et al., 2024; Henne et al., 2025; Herzke et al., 2023; Neuwald et al., 2022; Pickard et al., 2020; Pike et al., 2021; Scheurer et al., 2017; Scheurer & Nödler, 2021; Scott et al., 2005; Zheng et al., 2023).

As a PFAS, TFA falls within the scope of the proposed universal EU restriction on the substance class, which is motivated chiefly by its persistence. The toxicity of TFA is generally reported to be low across toxicology studies (Dekant & Dekant, 2023). Toxic effects in aquatic plants and algae arise in the range of 100s-1000s mg/L, which is orders of magnitude higher than current observed concentrations in aquatic environments (Garavagno et al., 2024; Hanson et al., 2024). The human margin of exposure was also reported by Dekant & Dekant (2023) to be much higher (4,000 - 476,000) than the standard of 100 set by the European Food Safety Authority (EFSA, 2023). Other authors critique these findings, stating a lack of ecological data and long-term studies that investigate chronic rather than acute toxicity (Arp et al., 2024; Hanson et al., 2024), and even misleading conclusions drawn by industry funded reports (Lysimachou et al., 2025). Despite TFA not being defined as bioaccumulative (ECHA, 2012) its persistency leads to external accumulation in the environment, including water and plants. This may lead to increased concentrations in the body due to intake of water and food that humans are exposed to (Arp et al., 2024). The German Federal Institute for Risk Assessment has also recently in May 2025 submitted an assessment to classify TFA as reprotoxic with the hazard statement “*May damage the unborn child. Suspected of impairing fertility*” (BfR, 2025) – noting, however, that the statement does not speak to health risks at current environmental exposure levels. This assessment was supported by the European Chemical Agency in June 2026 (*RAC 77 Minutes, Consensus on TFA Opinion on the Harmonised Classification for TFA*, 2026). In July 2026 the European Food Safety Authority lowered the acceptable daily intake to 0.014 mg/kg bodyweight per day, 3.5 times less than the previous limit, and set an acute reference dose of 0.07 mg/kg bodyweight (EFSA, 2026). It is furthermore less clear what the effects on vital Earth systems like biodiversity and biogeochemical cycles are. Arp et al. (2024) argue that TFA meets the criteria for a planetary boundary threat, based on criteria set up by (Persson et al., 2013) and further worked on by (MacLeod et al., 2014) which are stated as: disruptive effects on vital Earth ecosystems at a planetary scale that are poorly reversible. While the immediate negative impacts of TFA are not as profound as those of the more researched PFAS like PFOS or PFOA, its persistence, prevalence, and increasing presence in diverse matrices across the globe call for solutions to this massive contamination problem.

Removing TFA from a matrix like wastewater by conventional methods like ozonation and activated carbon filtration is ineffective (Ross et al., 2018; Scheurer et al., 2017), and alternatives like reverse osmosis are expensive, as well as water and energy intensive (Arp et al., 2024). An attractive alternative to these abiotic methods is microbial degradation of TFA.

PFAS have generally been considered immune to microbial breakdown (Hu & Scott, 2024; Skinner et al., 2025; Wackett, 2021). However, recent research provides hope that these are not truly ‘forever chemicals’ . The enzymatic capacity of the microbial world is vast, and enzymes may further act in a promiscuous manner on a wider range of compounds than their intended substrate. This facilitates the removal of pollutants through co-metabolism, rather than requiring the organisms to live purely off of the given compound - a proven strategy for degrading a range of other, recalcitrant pollutants (Hazen, 2010).

Enzymatic degradation of TFA poses a significant challenge. Despite being so common in the environment, fluorine is relatively absent from biology. This contrasts with other halogens: brominated, chlorinated, and iodinated compounds occur widely in nature, and the ability to degrade them is ancient. Industrial products containing these halogens have thus faced a ‘prepared and ready’ microbial population, accustomed to such highly halogenated compounds over millions of years. By contrast, very few fluorinated natural compounds exist (Chan & O’Hagan, 2012). Only a few enzymes capable of defluorination have been identified, degradation efficiencies vary, and degradation times can be weeks or months (Wackett, 2022). This paucity in natural fluorinated compounds is presented as a central factor in the rarity of microbes possessing enzymes capable of defluorination (Stockbridge & Wackett, 2024). Despite their paucity, naturally fluorinated compounds could be exactly the right starting point for further degradation.

Trifluoroacetate is the fully fluorinated form of acetate - but less fluorinated forms also exist, namely difluoroacetate (DFA) and (mono)fluoroacetate (MFA). In the context of microbial defluorination in general, and for TFA specifically, MFA is especially interesting to consider due to its analogous structure to TFA and the enzymes already evolved that can defluorinate it. Analogue enrichment is a strategy used to induce co- metabolic degradation of a specific compound, where a culture is enriched for degraders of a structural analogue. It has been shown to enrich microorganisms capable of co-metabolic degradation of other organic pollutants such as chlorophenol, aromatics, and polychlorinated biphenyls (De Los Cobos-Vasconcelos et al., 2006; Horvath, 1972; Leigh et al., 2006; Strubel et al., 1989).

MFA is one of the very few naturally occurring fluorine-containing compounds. It is a plant defence compound acting as a metabolic poison (Chan & O’Hagan, 2012) (Proudfoot et al., 2006). Being naturally evolved, the microbial world has had time to adapt to this compound. The ability to degrade MFA is well established in several, diverse microorganisms, including fungi and diverse genera of bacteria (Davis et al., 2012; Kelly, 1965; Seong et al., 2019; Twigg & Socha, 2001). Some are even able to utilize MFA as their only carbon source (Alexandrino et al., 2018; Twigg & Socha, 2001).

Goldman (1965) first described the enzymatic cleavage of the C-F bond in MFA. Since then, MFA has been shown to be degraded through hydrolytic cleavage of the C-F bond through a S_N_2-displacement reaction (Hu & Scott, 2024). Notable families of enzymes in this regard are fluoroacetate dehalogenases (FAcDs) and haloacid dehalogenases (HADs), which employ very similar mechanisms (Khusnutdinova et al., 2023) and function under aerobic conditions (Farajollahi et al., 2024). While often assumed to have evolved to detoxify and/or assimilate MFA (Stockbridge & Wackett, 2024) several fluoroacetate dehalogenases are somewhat promiscuous and have a wider substrate range. This can include cleaving other halogens (i.e. chloroacetate, bromoacetate, iodoacetate), or more ‘bulky’ and more fluorinated compounds (Hu & Scott, 2024). Interestingly, Khusnutdinova et al. (2023) screened eight purified enzymes with MFA-dehalogenating activity and found significant defluorination of DFA by three FAcDs. Although no successful TFA degradation was shown, Farajollahi et al. (2024) also showed defluorination of DFA by a FAcD and a HAD enzyme.

Overall, these findings encourage the theory that some, yet undiscovered, fluoroacetate dehalogenases could already possess the capacity to degrade TFA through promiscuous activity.

Reports of microbial degradation of TFA do exist, but are very few, are from the pre-genomics era, run over long timeframes, and do not identify any specific responsible microbial species or enzymes (Kim et al., 2000; Tang et al., 2024; Visscher et al., 1994). Visscher et al. (1994) applied small amounts (<1 μM) radiolabelled TFA to a salt marsh sediment slurry under both anoxic and oxic conditions. Results were obtained by HPLC analysis for ^14^C labelled TFA, DFA, MFA, acetate, and methane. The authors conclude trace levels of TFA can be degraded under both oxic and anoxic conditions, and that TFA is sequentially degraded to DFA, then MFA, and lastly to acetate with incubation times of 8-15 days. It should be noted that Emptage and colleagues (1997) attempted to replicate the result obtained by Visscher and colleagues without success. Kim et al. (2000) investigated degradation of TFA in an anaerobic chemostat reactor inoculated with an unspecified mixed culture of ethanol-degrading microorganisms. The authors report a complete degradation of TFA added to the bioreactor, which had a hydraulic retention time of 20 days.

Results were obtained by ion chromatography of F^-^ and LC analysis of TFA, DFA, and MFA. More recently Tang et al. (2024) incubated aquifer solids and its native microbiome with TFA and oxygen present and found a defluorination ratio of 3.46% (± 2.73%) after 10 months. F^-^ was measured using ion chromatography. While correlations between species abundance and F^-^ release were mapped, the study does not specify a concrete TFA degrading microorganism.

Other newer studies have not been able to demonstrate microbial or enzymatic TFA degradation. Alexandrino et al. (2018) reported complete defluorination of MFA by inocula from activated sludge, sediment, and rhizosphere samples after 21, 42 and 63 days, respectively. However, DFA and TFA proved wholly recalcitrant under all tested conditions. Results were obtained using a fluoride-ion selective electrode on sample supernatants. Farajollahi et al (2024) tested a HAD and a FAcD from *Delftia acidovorans* isolated from PFAS-contaminated soil. The enzyme showed activity for MFA and DFA, but none for TFA.

The state of the art presents a clear research gap: identifying microorganisms capable of degrading TFA as well as candidate catalytic enzymes.

The aim of the present study was to identify microorganisms capable of degrading TFA. Degraders were sought for through analogue enrichment with acetate, since TFA was hypothesized to be degraded cometabolically. Screening for degradation of TFA was performed with a fluoride ion-selective electrode (ISE), and degrading species were identified by Nanopore DNA sequencing, and candidate catabolic enzymes from genomic data were screened for with protein-ligand modelling. This was done by first setting up a sample of PFAS-contaminated wastewater sludge in media supplemented with acetate. Next, these cultures were exposed to MFA and screened for defluorination activity. When detecting F^-^ in a culture, an inoculum from it was used to set up new cultures incubated with DFA or TFA. The hypothesis was that organisms already possessing a fluoroacetate-degrading enzyme could have the ability to also degrade even more fluorinated versions of acetate. After obtaining candidate cultures they were all tested in a 30-day incubation experiment.

## Materials and methods

### Environmental sample

Sludge was provided by the Danish utility company Bornholms Energi & Forsyning from their plant in Boderne, Bornholm, Denmark. Here, the wastewater treatment plant pumps surplus sludge onto reed beds for mineralization. The wastewater primarily comes from a local town, Aakirkeby. Analyses of the sludge have shown significant PFAS pollution, to a degree where it cannot be used as agricultural fertilizer (not shown).

### Reagents

To make M9 minimal media a 5x M9 minimal salts solution was first prepared by mixing to a final concentration 0.239 M Na_2_HPO_4_•7H_2_O, 0.110 M KH_2_PO_4_, and 0.0935 M NH_4_Cl followed by autoclaving. The M9 minimal media itself was then prepared by mixing sterile reagents. A final concentration of 0.1 mM CaCl_2_ (1.0 M stock) and 2.0 mM MgSO_4_ (1.0 M stock) were added to MilliQ water along with 200 ml of 5x M9 minimal salts solution for a final volume of 1 L of M9 minimal media. Low-level TISAB was prepared as stated in the Thermo Scientific Orion Fluoride Ion Selective Electrode manual (Thermo Fisher Scientific, 2016). TFA, DFA, and MFA at >97% purity was purchased from Sigma-Aldrich, of which stock solutions were prepared in water to a concentration of 3 M. A 3 M acetate stock was prepared by mixing sodium acetate trihydrate with sterile MilliQ water and filter sterilizing the finished solution.

### Obtaining initial cultures / Enrichment and screening

Initial experiments were set up first to obtain microbial cultures from environmental samples to facilitate the screening for potential degraders. The first mesocosms were made by mixing 1 g of wastewater sludge and 3 ml M9 minimal media. The mesocosms were supplied with either 33 mM acetate, or 33 mM acetate with 33 mM MFA and incubated for 6 days. Subsequent cultures were set up using 100 µL of inoculum*μ* and varying concentrations of MFA, DFA, and TFA (17 - 66 mM) in 3 ml M9 minimal media. Cultures grown in M9 with 33 mM acetate were used as negative controls. Cultures were set up in Ø20 35 ml glass test tubes placed on a 200-rpm shaker to best allow for oxic conditions throughout the sample and incubated at 20 °C.

Acetate was chosen as the main carbon source by virtue of its analogous structure to TFA. The concentration of acetate supplied to samples was based on a 1:1 replacement of 0.2% glucose in terms of C-content. The utilised concentration range was then set to be similar to other experiments using acetate as a sole carbon source (Burckhardt et al., 2019; Kutscha & Pflügl, 2020; Oh et al., 2002; Petushkova & Tsygankov, 2017) to ensure a concentration high enough to supply adequate C to the culture for growth. To promote the likelihood of TFA degradation, the experiment used oxic conditions and high concentrations of TFA (Skinner et al., 2025), matching the acetate concentration. The relevant dehalogenase enzymes are expected to function under oxic conditions (Farajollahi et al., 2024).

### 30-day incubation experiment / Validation experiment

Efficiency of candidate degrading cultures was tested in a 30-day experiment with measurements of F^-^ at five timepoints (day 2, 5, 9, 15 and 30). All cultures were set up in triplicates in 4.5 ml M9 in Ø20 35 ml glass test tubes, each incubated with 33 mM acetate and either 33 mM MFA, DFA or TFA. Cultures were incubated at 28 °C and were placed on shakers at 200 rpm. Negative controls with cells growing on 33 mM acetate in M9 minimal media were set up in triplicate. Positive controls were made when measuring. To make positive controls 1 mM F^-^ in low-level TISAB was spiked in samples from the negative control mesocosm to a final concentration of 15 μM in the 400 μL. Sterile controls with 33 mM acetate and 33 mM of MFA, DFA, and TFA, respectively, were set up in singles.

### Fluoride measurements with fluoride selective electrode

To measure release of F^-^, a Thermo Scientific Orion Star A214 pH/ISE Meter and Orion fluoride selective electrode (ISE) was used on supernatants extracted from the cultures. To avoid the need for large culture volumes in a setup containing many samples, a 1.75x dilution (400 μL sample in 700 μL M9) was applied when measuring fluoride. First, 400 µL of sample was taken from each culture and centrifuged at 10,000 × g to pellet cells. The supernatant was transferred to a 15 ml falcon tube containing 700 µL M9 minimal media and 1.1 ml of low-level TISAB, giving a 1:1 relationship between low-level TISAB and sample matrix, as per instrument protocol. Low-level TISAB is appropriate for samples with a fluoride concentration up to 21 µM and no complexing agents like iron or aluminum. The limit of detection is 1 μM F^-^.

A standard curve was prepared for the purpose of quantifying F^-^ release (see Supplementary Fig. S1). To account for potential matrix effects of the media the standard curve was prepared by standard addition to M9. First, 5 mL of M9 was mixed with 5 mL of low-level TISAB. The sample was then spiked with 1 mM F^-^ in low-level TISAB in accordance with the instrument protocol provided by ThermoFisher (Thermo Fisher Scientific, 2016) adjusted to a total volume of 10 mL instead of 100 mL.

### 19F-NMR for fluoride and fluorinated metabolites

The ^19^F NMR spectra were acquired on an 18.8 T Bruker AVANCE IIIHD spectrometer operating at a ^19^F Larmor frequency of ν_19F_ = 752.74 MHz and equipped with a 5 mm TCI H/F CryoProbe (Bruker, Germany). Spectra were recorded using the standard pulse sequence *zgflqn* (Bruker), employing an 11ms p/2 excitation pulse, a relaxation delay of 8.5 seconds, an acquisition time of 1.46 seconds and 1000 transient scans. These parameters resulted in a total recycle time of roughly 10 seconds. Separate reference experiments on TFA and free fluoride demonstrated that this recycle time ensured recovery of >98% of the equilibrium magnetization. All spectra were measured at 25 °C. NMR samples were prepared by adding 60 μL of D2O (99.98%, Deutero GmBH, DE) to 120 μL of sample mixture, followed by transfer to a 3 mm NMR tube.

### DNA extraction and sequencing

DNA was extracted from presumed degraders using a Bead-Beat Micro AX Gravity (A&A Biotechnology, Gdansk, Poland). DNA purity was assessed using a NanoDrop ND-1000 Spectrophotometer (Thermo Fisher, MA, USA) on 1.5 μL of sample using the elution medium as a blank, and DNA was treated with a Zymo DNA Clean & Concentrator^TM^-5 kit where necessary. DNA concentration was determined by an Invitrogen Qubit^®^ 2.0 fluorometer (Thermo Fisher, MA, USA) using a 1X dsDNA High Sensitivity Assay kit on 2 μL of sample.

Sequencing of extracted DNA was performed by Nanopore sequencing. The extracted DNA was first prepared using a Rapid Barcoding Kit 24 (SQK-RBK114.24), yielding barcoded DNA libraries. The DNA libraries were loaded onto an Oxford Nanopore Technologies PromethION P2 Solo sequencer with an R10.4.1 flowcell. Basecalling was performed with dorado, as implemented in MinKnow v25.09.16.

Sequencing adapters were trimmed from raw reads with porechop v0.2.4 (Wick, 2017/2026) before genome assembly with Flye v2.9.5 (Kolmogorov et al., 2019). Assembled genomes were assessed for completeness with CheckM2 v1.2.2 (Parks et al., 2015) and contigs from multi-bacteria samples (metagenomes) were binned with SemiBin2 v2.2.1 (Pan et al., 2023). Binned metagenomes were assessed again with CheckM2. Taxonomy of binned contigs and complete genomes was assigned with GTDB-tk v2.6.1 (Chaumeil et al., 2020) and gene annotation was performed with Bakta v1.8.2 (Schwengers et al., 2021). In order to find candidate TFA-catalytic hydrolases, a sequence file consisting of 12 alpha/beta fold hydrolases (accession numbers CAE26606.1, AAG04199.1, ABE42492.1, BAE94252.1, CAD15064.1, AAZ48563.1, BAB77563.1, PZP66635.1, CAA39254.1, QAX87819.1, BAB85577.1, ABG92066.1) was used as query in a BLAST v2.17.0+ (Altschul et al., 1990) search against the assembled and binned metagenome of sample MFA1C. Co-folding of TFA and identified alpha/fold hydrolases was performed with Boltz-2.

### Statistical analysis

A two-way analysis of variance (ANOVA) mixed model was fitted to the data. Positive controls, sterile controls, and data for the calibration curve (Supplementary Table S1) were excluded from the ANOVA as these are not of statistical relevance. The model was fitted using lmer from the lmerTest package (Kuznetsova et al., 2017). The fitted model was analysed using the emmeans (Lenth & Piaskowski, 2017) and multcomp (Hothorn et al., 2008) packages. Model assumptions of normality and residual variance homogeneity were visually assessed using residual and QQ-plots (Supplementary Figs. S2 and S3). All statistical analysis was performed using R version 4.5.0 (R Core Team, 2025) in RStudio version 2026.1.0.392 (Posit team, 2026).

## Results and discussion

### All microbial consortia degrade MFA

Measures of defluorination of MFA, DFA, and TFA over time by the candidate cultures can be seen in figure 1A-C. For expanded experimental results see Supplementary Table S2. Samples have been named after the most fluorinated compound that the given culture was able to degrade before being selected for the main experiment. Defluorination of MFA occurred in all samples, and to a greater degree than DFA and TFA. This finding aligns with other studies on FAcD defluorination of MFA and DFA (Farajollahi et al., 2024; Jansen et al., 2026; Khusnutdinova et al., 2023), and with the fact that MFA can be fully metabolized. Curiously, triplicates diverged much more on MFA compared to DFA and TFA (Fig. 1). Stochastic effects of inoculation could be a relevant factor contributing to this result, especially in regard to presence or absence of rare species (Hayashi et al., 2026). While 100 µL of inoculum contains a large number of cells and should constitute a representative sample of the population, some biological variation in triplicates is still expected. If multiple degraders of MFA are present, but are relatively rare sub-groups in the population, differences in community composition in each sample could more easily arise, thus affecting degradation outcomes. The fact that MFA can be metabolized rather than co-metabolized could also facilitate more varied degradation outcomes. Since MFA is more degradable than DFA and TFA, samples will diverge faster and to a greater degree compared to samples where there is less activity overall. This in turn means that fluoride toxicity could also play a comparatively larger role in MFA samples. The toxicity of MFA itself could also pose an immediate selective pressure for its degradation (Jansen et al., 2026). All of these factors put together create a set of more complex conditions in MFA samples, which could extrapolate any initial stochasticity.

**Figure 1.**
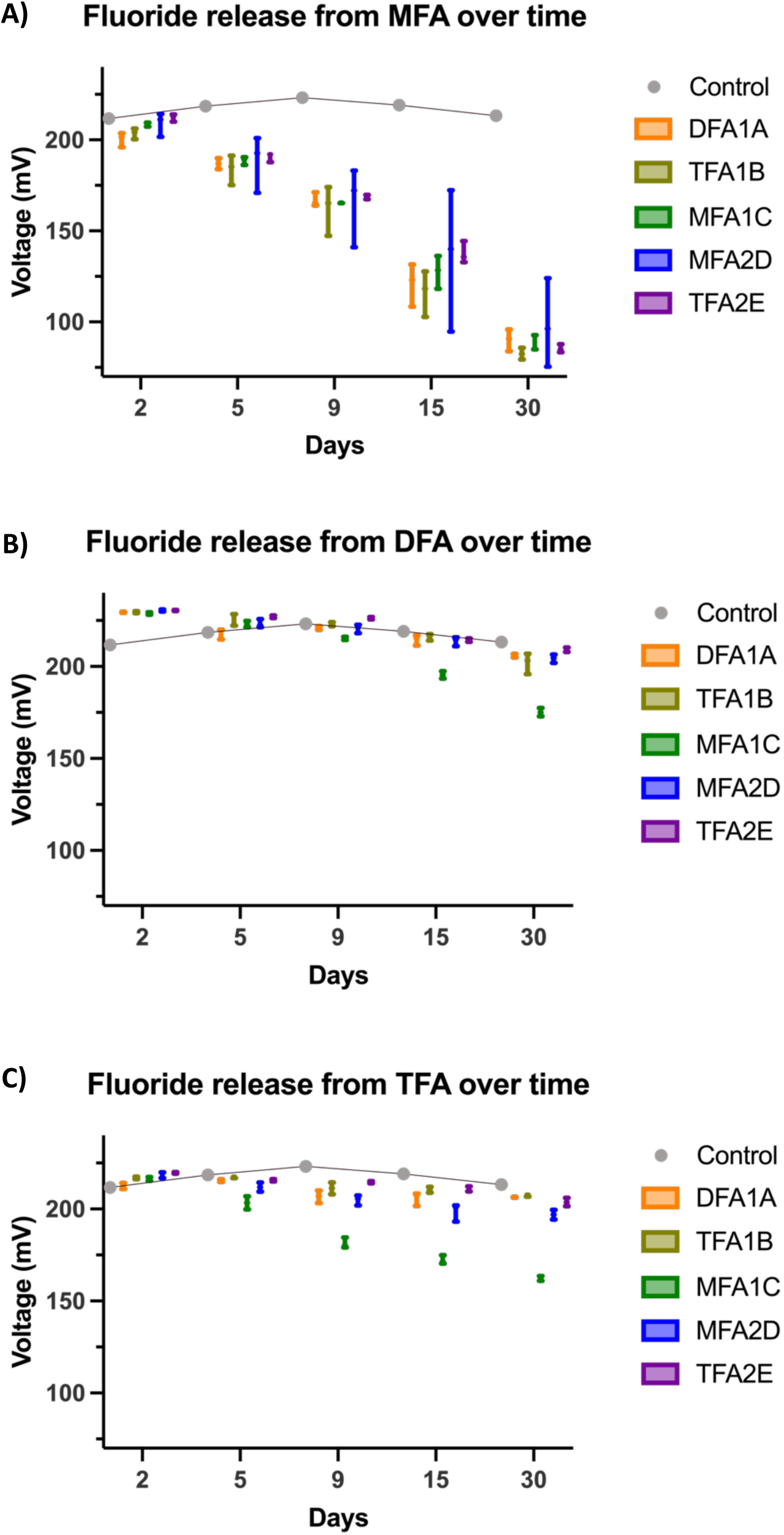
Voltage measurements of triplicate samples over 30 days as a measure of fluoride release form cultures incubated with either 33 mM MFA (A), DFA (B), or TFA (C). A lower voltage reading is associated with greater F^-^ release. The control sample is a negative control incubated only with 33 mM acetate. Cultures are partially named to indicate the fluorinated compound they were found to degrade in the discovery phase.

### One consortium degrades both DFA and TFA

Notably, panel C in figure 1 also shows the degradation capacity of a sample, MFA1C, with clear indications of fluoride release when incubated with both DFA and TFA. The sterile, abiotic controls did not exhibit defluorination, ruling out abiotic degradation in any culture (Supplementary Table S2). The defluorination capacity of sample MFA1C on DFA and TFA is greatest at the initial stages of incubation from day 0-9, whereafter it flattens out (Supplementary Fig. S4).

Figure 2 provides an overview of the two-way ANOVA mixed model test, showing total defluorination at the end of the 30-day experiment. The statistical analysis underscores the defluorination potential of culture MFA1C, which released significantly more fluoride than all other samples from both TFA and DFA (significance letter a, b, respectively). The TFA degradation in this best performing culture is estimated to be 11.2 ± 0.7 µM. This constitutes approximately 0.01% of the total fluoride from the 33 mM TFA applied (for full calculation see Supplementary Table S3).

**Figure 2.**
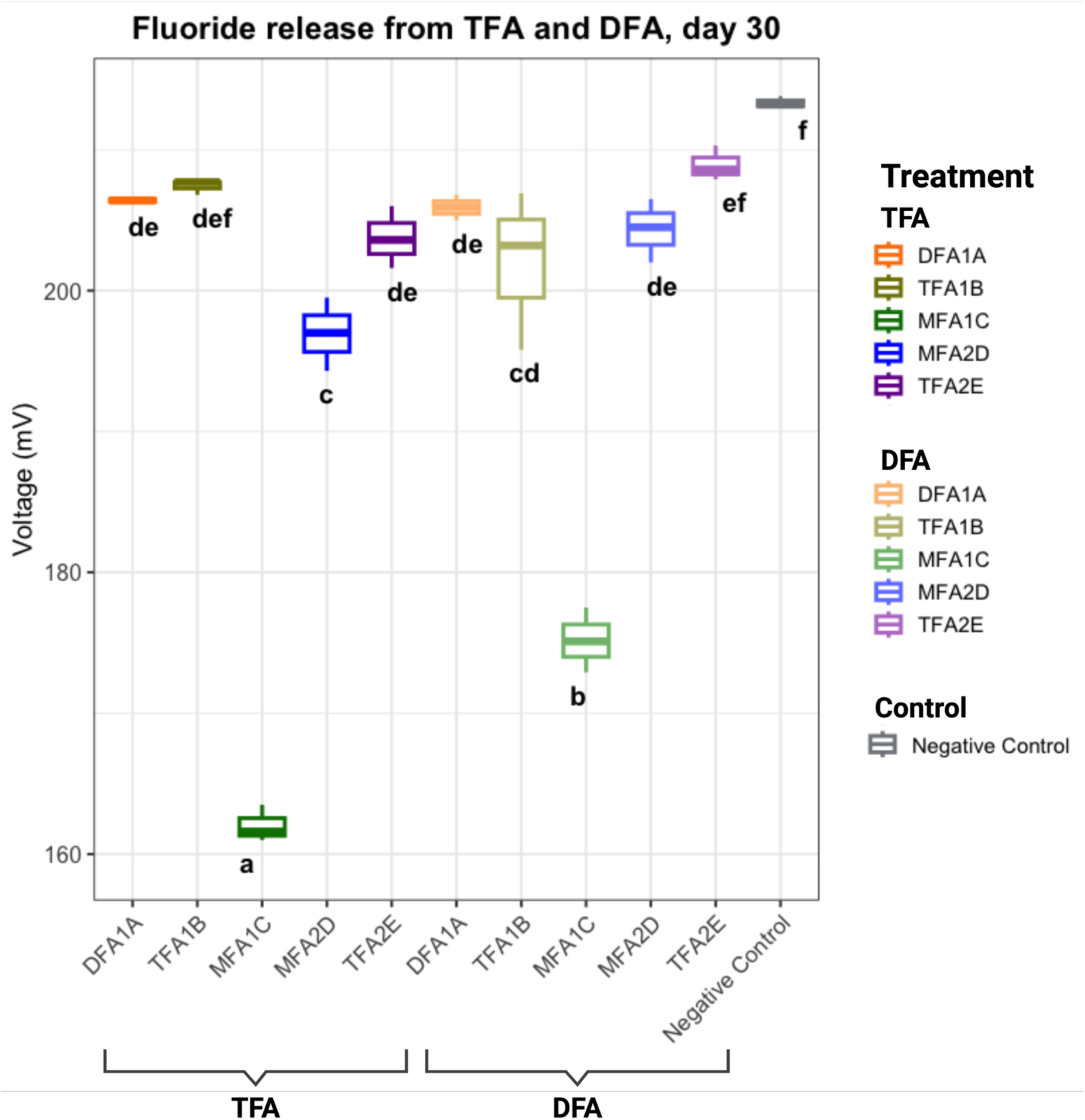
Comparison of culture group means on incubation day 30. Brackets below cultures indicate the compound supplied to the cultures. Statistical significance has been analysed using a two-way ANOVA mixed model at a 0.95-confidence level across total incubation time. The negative control was supplied only with acetate. Statistically significant differences between group means on day 30 are given by assignment of different letters.

There is some variation in electrode response to the negative sample which could likely be attributed to the instrument’s variability described previously. Most samples (DFA1A, TFA1B, TFA2E) do not differ markedly on day 30 from the negative control when looking at the total trajectory. Meaning, on day 30 the values might be slightly lower than the control on day 30, but not much different from the control on day 1. High mV of 200 or more (i.e. very low F^-^) in samples should be interpreted cautiously, as the electrode is most uncertain at low concentrations, in our experience.

### Orthogonal fluoride detection with ^19^F-NMR shows TFA degrading to DFA, MFA, and fluoride

To obtain compound-specific confirmation of the fluoride release measured by ISE, and to detect partially defluorinated intermediates, supernatants from the TFA-incubated cultures were analysed by ¹⁹F-NMR (Fig. 3). In all four cultures, the spectra were dominated by the TFA singlet at -75 ppm, and quantification against an external TFA calibration curve returned 32-34 mM, i.e. indistinguishable from the 33 mM supplied at the start of the experiment. In culture MFA1C, however, three further resonances were resolved that were absent from all other cultures: a doublet at −124.5 ppm and a triplet at -217 ppm, matching the chemical shifts and coupling patterns of the DFA and MFA reference compounds, respectively, and a broad signal at −120 ppm corresponding to free fluoride. Quantification gave 0.004 mM DFA, 0.001 mM MFA, and 0.006 mM F⁻ in this culture, corresponding to relative ¹⁹F integrals of 1.0:0.2:0.9. These proportions are internally consistent with stepwise hydrolytic defluorination: one C-F bond must be cleaved to form each DFA molecule and two to form each MFA molecule, which together predict 0.006 mM fluoride, in close agreement with the 0.006 mM measured directly. This is the same order of magnitude as, although somewhat lower than, the 11.2 ± 0.7 μM of TFA-derived fluoride estimated by ISE in the same culture. The detection of MFA further indicates that defluorination does not stop after the first C-F cleavage, but proceeds through DFA, in line with the ability of MFA1C to defluorinate DFA when supplied directly. Importantly, the TFA stock used to spike the cultures contained no detectable fluoride, DFA, or MFA when acquired with the same number of transients, excluding these species as impurities of the TFA source. The quantitative interpretation of these data is nevertheless limited: the degradation products occur at roughly four orders of magnitude lower concentration than the residual TFA and their signals have S/N < 20, so their absolute concentrations carry substantial uncertainty, while the corresponding depletion of TFA (ca. 0.015% of the pool) lies far below the ∼1.1% uncertainty of the TFA quantification. A closed fluorine mass balance therefore cannot be established from these measurements, and the ¹⁹F-NMR data are best read as direct, compound-specific evidence that MFA1C converts TFA to DFA, MFA, and fluoride, rather than as a quantitative measure of turnover.

**Figure 3.**
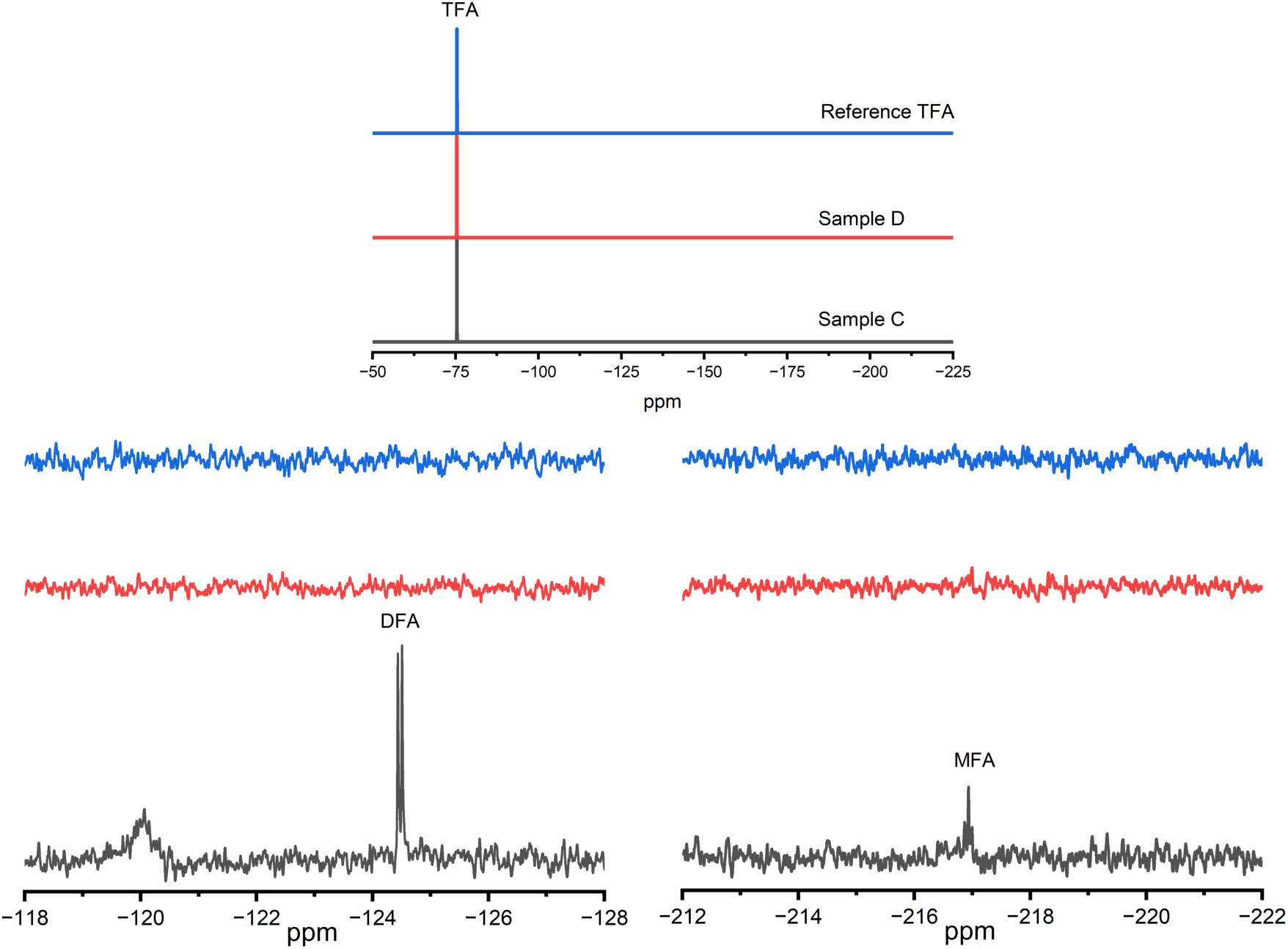
¹⁹F-NMR spectra of culture supernatants after 30 days of incubation with 33 mM TFA, shown with a spectrum of the TFA stock used to spike the cultures. The upper panel shows the full spectral range for the TFA reference (blue), culture MFA2D (red), and culture MFA1C (grey). These two samples are shown, as they represent a positive for defluorination of TFA (MFA1C) and a negative (MFA2D). All samples were run similarly on ^19^F-NMR. All three are dominated by the TFA singlet at -75 ppm. The lower panels are vertical expansions of the two regions containing defluorination products: −118 to −128 ppm (left), with the broad free fluoride signal at −120 ppm and the DFA doublet at −124.5 ppm, and -212 to -222 ppm (right), with the MFA triplet at -217 ppm. Fluoride, DFA, and MFA are present only in culture MFA1C and are absent from both culture MFA2D and the TFA reference. Note that the product signals are approximately four orders of magnitude smaller than the TFA signal, and that the vertical scale therefore differs between the upper and lower panels. All spectra were acquired with 1,000 transients, and assignments were verified against reference spectra of TFA, DFA, and MFA (Supplementary Fig. S5).

### DNA sequencing and protein ligand modelling yields two candidate enzymes

DNA sequencing identified a range of bacteria present in cultures supplied TFA and DFA; three species of *Achromobacter*, an *Advenella* sp., and two species of *Brucella*, *Serratia, Sphingobacterium,* and *Stenotrophomonas*. Some genomes were only identified to the genus level. Sequencing of sample TFA2E grown with TFA unfortunately did not yield any sequencing data but did also not release measurable fluoride on TFA (Fig. 1). All identified genera consist of aerobic, Gram-negative bacteria, aligning with the oxic experimental conditions. No genera found in the TFA experiment performed by Tang et al. (2024) were identified in this study. Despite being inoculated from the same culture, there are generally differences in the sequenced species in the DFA vs. TFA treatment; the same species were not found in DFA1A incubated with TFA and with DFA, for example. This could be due to the stochastic effects discussed previously, resulting in some species being absent or in too low concentrations to be accurately sequenced. Protein-ligand co-folding of enzymes in the MFA1C sequenced genomes identified two alpha/beta fold hydrolase candidate enzymes from *Achromobacter denitrificans* and *Stenotrophomonas* sp. identified at genus level. These enzymes are subject to further, ongoing testing both *in silico* and *in vivo*.

**Table 1.**
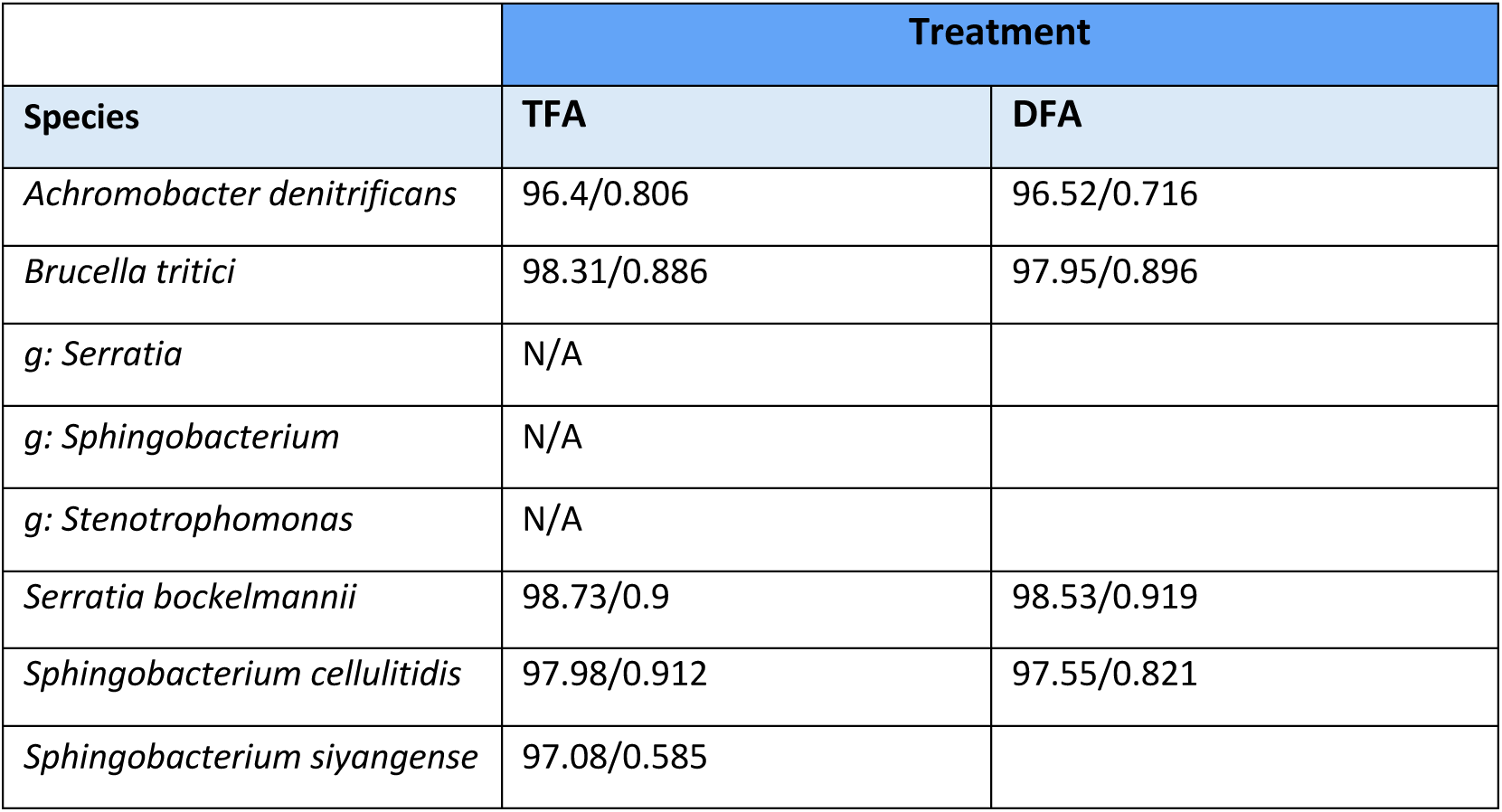
Overview of sequencing results from sample MFA1C in the main experiment. Numbers indicate closest genome Average Nucleotide Identity (ANI) / closest genome Aligned Fraction (AF). ANI values are a measure of % genome resemblance on a nucleotide level of comparable sequences between the sequenced DNA and genomic database species sequences. AF values indicate the ratio of the genome that could be compared between sequenced DNA and the database sequences

|  | Treatment |  |
| --- | --- | --- |
| Species | TFA | DFA |
| *Achromobacter denitrificans* | 96.4/0.806 | 96.52/0.716 |
| *Brucella tritici* | 98.31/0.886 | 97.95/0.896 |
| *g: Serratia* | N/A |  |
| *g: Sphingobacterium* | N/A |  |
| *g: Stenotrophomonas* | N/A |  |
| *Serratia bockelmannii* | 98.73/0.9 | 98.53/0.919 |
| *Sphingobacterium cellulitidis* | 97.98/0.912 | 97.55/0.821 |
| *Sphingobacterium siyangense* | 97.08/0.585 |  |

### Stability of microbial consortia

The initial expectation going into the main experiment was that cultures where TFA degradation had already occurred would continue to defluorinate TFA and possibly be the best performing ones. Counter to this expectation, TFA1B did not degrade TFA at all, and TFA2E only did so poorly. Instead, the two most promising degraders were MFA1C and MFA2D. Out of the two, MFA2D had been the most efficient MFA degrader, but was outperformed by MFA1C for TFA degradation. However, as was also observed by Farajollahi et al. (2024), the better MFA degrading enzyme was in their case not the better DFA degrader. Thus, degradation efficiency of one compound alone does not seem an accurate proxy for the efficiency on other compounds. The occurrence of TFA degradation by cultures able to degrade MFA supports the hypothesis of co-metabolic degradation by promiscuous enzymes. Cultures were not tested systematically for growth on pure fluorinated compounds, but small tests concurrent with the development of the main experiment showed no growth on either DFA or TFA when provided without acetate (not shown). This further supports that the degradation observed is happening via co-metabolism.

### Challenges of TFA defluorination

The application of microorganisms to degrade a PFAS like TFA faces some specific and significant challenges due to the chemical nature of the compound, chiefly the inertness and strength of the C-F bond (Khan, 2025) and the toxicity of F^-^ (McIlwain et al., 2021). A TFA degrader must thus possess traits to pass several hurdles at once as outlined by Wackett (2021): TFA must be transported across the cell membrane (in the case of alpha/beta fold hydrolases that are intracellular), TFA must be degraded enzymatically, and intracellular F^-^ must be excreted. Considering this combination of challenges, the capacity to degrade TFA was expected to be a rare trait. Aligning with this, the ability to degrade MFA only sometimes translated to a capacity for TFA degradation in our consortia. Low degradation rates were also expected due to the hypothesis of co-metabolic degradation and general compound recalcitrance. Besides those factors influencing degradation capacity, potential enzyme inhibition and perhaps F^-^ toxicity could play a further role.

An important factor in culture performance and stability could be negative selective pressures associated with fluoride toxicity. Fluorine is quite a common element - the 13^th^ most abundant in the Earth’s crust. As such, fluoride leached from minerals or dispersed through volcanic activity reach environmental concentrations of around 30-80 µM in oceans and 10-400 µM in surface and ground water (McIlwain et al., 2021; Stockbridge & Wackett, 2024). At an intracellular pH of 7-7.5 fluorine will exist mainly as F^-^ (HF p*K*_a_ = 3.4), which cannot readily diffuse out of the cell again. Here, F^-^ inhibits a broad spectrum of essential enzymes (McIlwain et al., 2021). This impairment posed by F-toxicity demands a way for microorganisms to defend themselves. Species across life domains possess resistance mechanisms, namely genetic switches (riboswitches) sensitive and specific to F^-^ that regulate gene expression for its export and for mitigation of toxic effects (Baker et al., 2012). The F^-^ inhibitory constant for enzymes is around 100 µM. Cells start defending themselves by changing gene expression at around 60 µM intracellular and 200 µM extracellular concentrations (Stockbridge & Wackett, 2024). The 11.2 µM reported here falls below these limits at first glance. However, with a typical bacterial cell volume of 10^-9^ µL concentrations of F^-^ can become significant (Wackett, 2022). In this example it would take around 36,000 F^-^ atoms to create an intracellular concentration of 60 µM, or roughly 12,000 TFA molecules. If intracellular fluoride concentrations rise too quickly for cells to cope by exporting fluoride, they could either be inhibited or killed, although we do not believe that the current slow TFA defluorination would lead to such inhibitory concentrations.

Ideally, evolving the capacity to degrade a new substrate confers a competitive advantage in the form of access to energy, carbon, or nutrients (i.e. N or P). The toxicity of fluoride released intracellularly, however, imposes a negative selection pressure on any successful degraders. Combined with the lack of positive selection due to the recalcitrance, lack of nutrients, and oxidized nature of TFA, any degradation likely only hampers the responsible organism (Wackett, 2024). The decline of degradation efficiency over time, and the somewhat transient nature of degradation capability across reinoculations, could likely be due to these disadvantages and lack of advantages associated with TFA defluorination. Furthermore, a long form incubation experiment as described here does not provide optimized growth conditions in the form of fresh media, a pH adjusted and buffered environment, and resupply of nutrients. Based on our experimental data and the general knowledge of metabolism of fluorinated compounds, we believe that TFA degradation is rare, allocated to co-metabolism, and generally under negative selective pressure. Other studies of MFA and DFA degradation have also noted a decrease in enzymatic activity over time. Results from Jansen et al. (2026) indicated that FAcDs were covalently modified and thereby inactivated when degrading DFA.

Permanent enzymatic inhibition could then be another plausible explanation for the decline in defluorination rate over time.

We also expected that DFA would be more easily degraded than TFA by virtue of being less fluorinated. Interestingly, counter to that expectation, the efficiency in degrading TFA was statistically higher than the degradation of DFA. It could be the case that degrading enzymes experience differential inhibition by the degradation products of each compound. It would seem the process of catalysing TFA, presumably to DFA, poses less enzymatic inhibition than the presumed catalysis of DFA to MFA.

### Outlook

The experimental framework of analogue enrichment with acetate proved effective in promoting TFA degrading cultures. To the best of our knowledge, the work presented herein is the first to identify specific organisms and candidate enzymes capable of degrading TFA. We are currently attempting to get single isolates from the consortium and are performing heterologous expression of candidate alpha/beta fold hydrolase enzymes.

A natural next step is investigating the enzymatic mechanism and kinetics. Testing the function of candidate enzymes by cloning and heterologous expression is essential. The documentation of TFA degradation and identification of candidate TFA degrading enzymes presented in this work ultimately facilitates the pursuit of bioremediation of TFA. Bioremediation generally faces significant challenges for *in situ* applications, namely poor survivability and effectiveness of the applied microbiome in bioaugmentation (Megharaj et al., 2011). However, in light of the complete paucity of known TFA degraders and associated enzymes until now, the identification of such an organism here is a keystone finding. A promiscuous enzyme with capacity to degrade a PFAS like TFA promotes new avenues of research for tackling PFAS pollution, and work towards biological treatment of TFA pollution as a cheap, environmentally friendly solution.

## Acknowledgements.

The authors would like to thank project leader Torben Jørgensen from Bornholms Energi & Forsyning for providing the environmental samples.

This project was supported by Independent Research Fund Denmark [grant numbers 3105-00303B and 5254-00090B awarded to TKN].

## References

1. Alexandrino, D. A. M., Ribeiro, I., Pinto, L. M., Cambra, R., Oliveira, R. S., Pereira, F., & Carvalho, M. F. (2018). Biodegradation of mono-, di- and trifluoroacetate by microbial cultures with different origins. *New Biotechnology*, International Society of Environmental Biotechnology 2016, *43*, 23–29. 10.1016/j.nbt.2017.08.005

2. Altschul, S. F., Gish, W., Miller, W., Myers, E. W., & Lipman, D. J. (1990). Basic local alignment search tool. Journal of Molecular Biology, 215(3), 403–410. 10.1016/S0022-2836(05)80360-2

3. Arp, H. P. H., Gredelj, A., Glüge, J., Scheringer, M., & Cousins, I. T. (2024). The Global Threat from the Irreversible Accumulation of Trifluoroacetic Acid (TFA). Environmental Science & Technology, 58(45), 19925–19935. 10.1021/acs.est.4c06189

4. Baker, J. L., Sudarsan, N., Weinberg, Z., Roth, A., Stockbridge, R. B., & Breaker, R. R. (2012). Widespread Genetic Switches and Toxicity Resistance Proteins for Fluoride. Science, 335(6065), 233–235. 10.1126/science.1215063

5. Berhanu, A., Mutanda, I., Taolin, J., Qaria, M. A., Yang, B., & Zhu, D. (2023). A review of microbial degradation of per- and polyfluoroalkyl substances (PFAS): Biotransformation routes and enzymes. Science of The Total Environment, 859, 160010. 10.1016/j.scitotenv.2022.160010

6. BfR, G. F. I. for R. A. (2025, May 26). Trifluoroacetic acid (TFA): Assessment for classification in new hazard classes submitted. Bundesinstitut Für Risikobewertung. https://www.bfr.bund.de/en/press-release/trifluoroacetic-acid-tfa-assessment-for-classification-in-new-hazard-classes-submitted/

7. Buck, R. C., Franklin, J., Berger, U., Conder, J. M., Cousins, I. T., de Voogt, P., Jensen, A. A., Kannan, K., Mabury, S. A., & van Leeuwen, S. P. (2011). Perfluoroalkyl and Polyfluoroalkyl Substances in the Environment: Terminology, Classification, and Origins. Integrated Environmental Assessment and Management, 7(4), 513–541. 10.1002/ieam.258

8. Burckhardt, R. M., Buckner, B. A., & Escalante-Semerena, J. C. (2019). Staphylococcus aureus modulates the activity of acetyl-Coenzyme A synthetase (Acs) by sirtuin-dependent reversible lysine acetylation. Molecular Microbiology, 112(2), 588–604. 10.1111/mmi.14276

9. Chan, K. K. J., & O’Hagan, D. (2012). The Rare Fluorinated Natural Products and Biotechnological Prospects for Fluorine Enzymology. In Methods in Enzymology (Vol. 516, pp. 219–235). Academic Press. 10.1016/B978-0-12-394291-3.00003-4

10. Chaumeil, P.-A., Mussig, A. J., Hugenholtz, P., & Parks, D. H. (2020). GTDB-Tk: A toolkit to classify genomes with the Genome Taxonomy Database. Bioinformatics, 36(6), 1925–1927. 10.1093/bioinformatics/btz848

11. Davis, C. K., Webb, R. I., Sly, L. I., Denman, S. E., & McSweeney, C. S. (2012). Isolation and survey of novel fluoroacetate-degrading bacteria belonging to the phylum Synergistetes. FEMS Microbiology Ecology, 80(3), 671–684. 10.1111/j.1574-6941.2012.01338.x

12. De Los Cobos-Vasconcelos, D., Santoyo-Tepole, F., Juárez-Ramírez, C., Ruiz-Ordaz, N., & Galíndez-Mayer, C. J. J. (2006). Cometabolic degradation of chlorophenols by a strain of *Burkholderia* in fed-batch culture. *Enzyme and Microbial Technology*, Papers from the 1st International Conference on Environmental, Industrial and Applied Microbiology (BioMicroWorld-2005), 40(1), 57–60. 10.1016/j.enzmictec.2005.10.038

13. Dekant, W., & Dekant, R. (2023). Mammalian toxicity of trifluoroacetate and assessment of human health risks due to environmental exposures. Archives of Toxicology, 97(4), 1069–1077. 10.1007/s00204-023-03454-y

14. DeWitt, J. C., Goldenman, G., Lohmann, R., Ng, C. A., & Wang, Z. (2025). Emerging toxicological awareness of per- and polyfluoroalkyl substances: The rising concern over ‘forever chemicals’. Disease Models & Mechanisms, 18(11), dmm052647. 10.1242/dmm.052647

15. Duan, Y., Sun, H., Yao, Y., Meng, Y., & Li, Y. (2020). Distribution of novel and legacy per-/polyfluoroalkyl substances in serum and its associations with two glycemic biomarkers among Chinese adult men and women with normal blood glucose levels. Environment International, 134, 105295. 10.1016/j.envint.2019.105295

16. ECHA, E. C. A. (2012). *Trifluoroacetic acid 100.000.846 | 512f4572-7d1e-43de-8009-2632def5062a—ECHA CHEM*. https://chem.echa.europa.eu/100.000.846/dossier-view/512f4572-7d1e-43de-8009-2632def5062a/e08b7134-3f0d-4351-ab3f-51b48497b756_125ed25b-eae2-4036-a459-db4c0c66f231?searchText=76-05-1

17. EFSA, E. F. S. A. (2023, April 27). *Margin of Exposure | EFSA*. https://www.efsa.europa.eu/en/topics/topic/margin-exposure

18. EFSA (European Food Safety Authority). (2026). Scientific report on consumer health-based guidance values for trifluoroacetic acid. EFSA Journal, 24(7), e10227. 10.2903/j.efsa.2026.10227

19. Emptage, M., Tabinowski, J., & Odom, J. M. (1997). Effect of Fluoroacetates on Methanogenesis in Samples from Selected Methanogenic Environments. Environmental Science & Technology, 31(3), 732–734. 10.1021/es9603822

20. European Commission. (2017). *Residues of DFA and TFA in Samples of Plant Origin* [EURL-SRM report]. EU Reference Laboratories for Residues of Pesticides. https://www.eurl-pesticides.eu/userfiles/file/eurlsrm/eurlsrm_residue-observation_tfa-dfa.pdf

21. Farajollahi, S., Lombardo, N. V., Crenshaw, M. D., Guo, H.-B., Doherty, M. E., Davison, T. R., Steel, J. J., Almand, E. A., Varaljay, V. A., Suei-Hung, C., Mirau, P. A., Berry, R. J., Kelley-Loughnane, N., & Dennis, P. B. (2024). Defluorination of Organofluorine Compounds Using Dehalogenase Enzymes from *Delftia acidovorans* (D4B). ACS Omega, 9(26), 28546–28555. 10.1021/acsomega.4c02517

22. Freeling, F., & Björnsdotter, M. K. (2023). Assessing the environmental occurrence of the anthropogenic contaminant trifluoroacetic acid (TFA). Current Opinion in Green and Sustainable Chemistry, 41, 100807. 10.1016/j.cogsc.2023.100807

23. Freeling, F., Scheurer, M., Koschorreck, J., Hoffmann, G., Ternes, T. A., & Nödler, K. (2022). Levels and Temporal Trends of Trifluoroacetate (TFA) in Archived Plants: Evidence for Increasing Emissions of Gaseous TFA Precursors over the Last Decades. Environmental Science & Technology Letters, 9(5), 400–405. 10.1021/acs.estlett.2c00164

24. Garavagno, M. de los A., Holland, R., Khan, M. A. H., Orr-Ewing, A. J., & Shallcross, D. E. (2024). Trifluoroacetic Acid: Toxicity, Sources, Sinks and Future Prospects. Sustainability, 16(6), 2382. 10.3390/su16062382

25. Glüge, J., Scheringer, M., Cousins, I. T., DeWitt, J. C., Goldenman, G., Herzke, D., Lohmann, R., Ng, C. A., Trier, X., & Wang, Z. (2020). An overview of the uses of per- and polyfluoroalkyl substances (PFAS). Environmental Science: Processes & Impacts, 22(12), 2345–2373. 10.1039/D0EM00291G

26. Goldman, P. (1965). The Enzymatic Cleavage of the Carbon-Fluorine Bond in Fluoroacetate. Journal of Biological Chemistry, 240(8), 3434–3438. 10.1016/S0021-9258(18)97236-4

27. Hanson, M. L., Madronich, S., Solomon, K., Sulbaek Andersen, M. P., & Wallington, T. J. (2024). Trifluoroacetic Acid in the Environment: Consensus, Gaps, and Next Steps. Environmental Toxicology and Chemistry, 43(10), 2091–2093. 10.1002/etc.5963

28. Hayashi, I., Sánchez-Pinillos, M., & Toju, H. (2026). Stochastic Forces in Microbial Community Assembly: Founding Community Size Governs Divergent Ecological Trajectories. Ecology Letters, 29(5), e70388. 10.1111/ele.70388

29. Hazen, T. C. (2010). Cometabolic Bioremediation. In Handbook of Hydrocarbon and Lipid Microbiology (pp. 2505–2514). Springer, Berlin, Heidelberg. 10.1007/978-3-540-77587-4_185

30. Henne, S., Storck, F. R., Wöhrnschimmel, H., Leuenberger, M., Vollmer, M. K., & Reimann, S. (2025). Trifluoroacetate (TFA) in precipitation and surface waters in Switzerland: Trends, source attribution, and budget. Atmospheric Chemistry and Physics, 25(23), 18157–18186. 10.5194/acp-25-18157-2025

31. Herzke, D., Nikiforov, V., Yeung, L. W. Y., Moe, B., Routti, H., Nygård, T., Gabrielsen, Geir. W., & Hanssen, L. (2023). Targeted PFAS analyses and extractable organofluorine – Enhancing our understanding of the presence of unknown PFAS in Norwegian wildlife. Environment International, 171, 107640. 10.1016/j.envint.2022.107640

32. Horvath, R. S. (1972). Microbial Co-Metabolism and the Degradation of Organic Compounds in Nature. 36.

33. Hothorn, T., Bretz, F., & Westfall, P. (2008). Simultaneous Inference in General Parametric Models. Biometrical Journal, 50(3), 346--363.

34. Hu, M., & Scott, C. (2024). Toward the development of a molecular toolkit for the microbial remediation of per-and polyfluoroalkyl substances. Applied and Environmental Microbiology, 90(4), e00157–24. 10.1128/aem.00157-24

35. Jansen, S. C., van Beers, P., & Mayer, C. (2026). Engineering Fluoroacetate Dehalogenase by Growth-Based Selections on Non-Natural Organofluorides. Angewandte Chemie International Edition, 65(10), e24234. 10.1002/anie.202524234

36. Kelly, M. (1965). Isolation of Bacteria able to metabolize Fluoroacetate or Fluoroacetamide. Nature, 208(5012), 809–810. 10.1038/208809a0

37. Khan, M. F. (2025). Recent Progress and Challenges in Microbial Defluorination and Degradation for Sustainable Remediation of Fluorinated Xenobiotics. Processes, 13(7), 2017. 10.3390/pr13072017

38. Khusnutdinova, A. N., Batyrova, K. A., Brown, G., Fedorchuk, T., Chai, Y. S., Skarina, T., Flick, R., Petit, A.-P., Savchenko, A., Stogios, P., & Yakunin, A. F. (2023). Structural insights into hydrolytic defluorination of difluoroacetate by microbial fluoroacetate dehalogenases. The FEBS Journal, 290(20), 4966– 4983. 10.1111/febs.16903

39. Kim, B. R., Suidan, M. T., Wallington, T. J., & Du, X. (2000). Biodegradability of Trifluoroacetic Acid. Environmental Engineering Science, 17(6), 337–342. 10.1089/ees.2000.17.337

40. Kolmogorov, M., Yuan, J., Lin, Y., & Pevzner, P. A. (2019). Assembly of long, error-prone reads using repeat graphs. Nature Biotechnology, 37(5), 540–546. 10.1038/s41587-019-0072-8

41. Kutscha, R., & Pflügl, S. (2020). Microbial Upgrading of Acetate into Value-Added Products—Examining Microbial Diversity, Bioenergetic Constraints and Metabolic Engineering Approaches. International Journal of Molecular Sciences, 21(22), 8777. 10.3390/ijms21228777

42. Kuznetsova, A., Brockhoff, P. B., & Christensen, R. H. B. (2017). **lmerTest** Package: Tests in Linear Mixed Effects Models. Journal of Statistical Software, 82(13). 10.18637/jss.v082.i13

43. Leigh, M. B., Prouzová, P., Macková, M., Macek, T., Nagle, D. P., & Fletcher, J. S. (2006). Polychlorinated Biphenyl (PCB)-Degrading Bacteria Associated with Trees in a PCB-Contaminated Site. Applied and Environmental Microbiology, 72(4), 2331–2342. 10.1128/AEM.72.4.2331-2342.2006

44. Lenth, R. V., & Piaskowski, J. (2017). emmeans: Estimated Marginal Means, aka Least-Squares Means (p. 2.0.3) [Data set]. 10.32614/CRAN.package.emmeans

45. Lysimachou, A., Muilerman, H., & Roynel, S. (2025). *Manufacturing Doubt how industry downplays TFA’s toxicity*. Pesticide Action Network Europe (PAN Europe). https://www.pan-europe.info/sites/pan-europe.info/files/public/resources/reports/Manufacturing%20Doubt%20how%20industry%20downplays%20TFA%27s%20toxicity_Report%2026092025.pdf

46. MacLeod, M., Breitholtz, M., Cousins, I. T., Wit, C. A. de, Persson, L. M., Rudén, C., & McLachlan, M. S. (2014). Identifying Chemicals That Are Planetary Boundary Threats. Environmental Science & Technology, 48(19), 11057–11063. 10.1021/es501893m

47. McIlwain, B. C., Ruprecht, M. T., & Stockbridge, R. B. (2021). Membrane exporters of fluoride ion. Annual Review of Biochemistry, 90, 559–579. 10.1146/annurev-biochem-071520-112507

48. Megharaj, M., Ramakrishnan, B., Venkateswarlu, K., Sethunathan, N., & Naidu, R. (2011). Bioremediation approaches for organic pollutants: A critical perspective. Environment International, 37(8), 1362–1375. 10.1016/j.envint.2011.06.003

49. Mueller, R., & Schlosser, K. E. (2020). History and Use of Per- and Polyfluoroalkyl Substances (PFAS) found in the Environment.

50. Neuwald, I. J., Hübner, D., Wiegand, H. L., Valkov, V., Borchers, U., Nödler, K., Scheurer, M., Hale, S. E., Arp, H. P. H., & Zahn, D. (2022). Ultra-Short-Chain PFASs in the Sources of German Drinking Water: Prevalent, Overlooked, Difficult to Remove, and Unregulated. Environmental Science & Technology, 56(10), 6380–6390. 10.1021/acs.est.1c07949

51. Oh, M.-K., Rohlin, L., Kao, K. C., & Liao, J. C. (2002). Global Expression Profiling of Acetate-grown *Escherichia coli*∗. Journal of Biological Chemistry, 277(15), 13175–13183. 10.1074/jbc.M110809200

52. Pan, S., Zhao, X.-M., & Coelho, L. P. (2023). SemiBin2: Self-supervised contrastive learning leads to better MAGs for short- and long-read sequencing. Bioinformatics, 39(Supplement_1), i21–i29. 10.1093/bioinformatics/btad209

53. Parks, D. H., Imelfort, M., Skennerton, C. T., Hugenholtz, P., & Tyson, G. W. (2015). CheckM: Assessing the quality of microbial genomes recovered from isolates, single cells, and metagenomes. Genome Research, 25(7), 1043–1055. 10.1101/gr.186072.114

54. Persson, L. M., Breitholtz, M., Cousins, I. T., de Wit, C. A., MacLeod, M., & McLachlan, M. S. (2013). Confronting Unknown Planetary Boundary Threats from Chemical Pollution. Environmental Science & Technology, 47(22), 12619–12622. 10.1021/es402501c

55. Petushkova, E. P., & Tsygankov, A. A. (2017). Acetate metabolism in the purple non-sulfur bacterium Rhodobacter capsulatus. Biochemistry (Moscow), 82(5), 587–605. 10.1134/S0006297917050078

56. Pickard, H. M., Criscitiello, A. S., Persaud, D., Spencer, C., Muir, D. C. G., Lehnherr, I., Sharp, M. J., De Silva, A. O., & Young, C. J. (2020). Ice Core Record of Persistent Short-Chain Fluorinated Alkyl Acids: Evidence of the Impact From Global Environmental Regulations. Geophysical Research Letters, 47(10), e2020GL087535. 10.1029/2020GL087535

57. Pike, K. A., Edmiston, P. L., Morrison, J. J., & Faust, J. A. (2021). Correlation Analysis of Perfluoroalkyl Substances in Regional U.S. Precipitation Events. Water Research, 190, 116685. 10.1016/j.watres.2020.116685

58. Posit team. (2026). RStudio: Integrated Development Environment for R (Version 2026.1.0.392) [Computer software]. Posit Software, PBC. http://www.posit.co/

59. Proudfoot, A. T., Bradberry, S. M., & Vale, J. A. (2006). Sodium Fluoroacetate Poisoning.

60. R Core Team. (2025). R: A Language and Environment for Statistical Computing (Version 4.5.0) [Computer software]. R Foundation for Statistical Computing. https://www.R-project.org/

61. Richey, D. G., Driscoll, C. T., & Likens, G. E. (1997). Soil Retention of Trifluoroacetate. Environmental Science & Technology, 31(6), 1723–1727. 10.1021/es960649x

62. Ross, I., McDonough, J., Miles, J., Storch, P., Thelakkat Kochunarayanan, P., Kalve, E., Hurst, J., S. Dasgupta, S., & Burdick, J. (2018). A review of emerging technologies for remediation of PFASs. Remediation Journal, 28(2), 101–126. 10.1002/rem.21553

63. Scheurer, M., & Nödler, K. (2021). Ultrashort-chain perfluoroalkyl substance trifluoroacetate (TFA) in beer and tea – An unintended aqueous extraction. Food Chemistry, 351, 129304. 10.1016/j.foodchem.2021.129304

64. Scheurer, M., Nödler, K., Freeling, F., Janda, J., Happel, O., Riegel, M., Müller, U., Storck, F. R., Fleig, M., Lange, F. T., Brunsch, A., & Brauch, H.-J. (2017). Small, mobile, persistent: Trifluoroacetate in the water cycle – Overlooked sources, pathways, and consequences for drinking water supply. Water Research, 126, 460–471. 10.1016/j.watres.2017.09.045

65. Schwengers, O., Jelonek, L., Dieckmann, M. A., Beyvers, S., Blom, J., & Goesmann, A. (2021). Bakta: Rapid and standardized annotation of bacterial genomes via alignment-free sequence identification. Microbial Genomics, 7(11), 000685. 10.1099/mgen.0.000685

66. Scott, B. F., Macdonald, R. W., Kannan, K., Fisk, A., Witter, A., Yamashita, N., Durham, L., Spencer, C., & Muir, D. C. G. (2005). Trifluoroacetate Profiles in the Arctic, Atlantic, and Pacific Oceans. Environmental Science & Technology, 39(17), 6555–6560. 10.1021/es047975u

67. Seong, H. J., Kwon, S. W., Seo, D.-C., Kim, J.-H., & Jang, Y.-S. (2019). Enzymatic defluorination of fluorinated compounds. Applied Biological Chemistry, 62(1), 62. 10.1186/s13765-019-0469-6

68. Skinner, J. P., Raderstorf, A., Rittmann, B. E., & Delgado, A. G. (2025). Biotransforming the “Forever Chemicals”: Trends and Insights from Microbiological Studies on PFAS. Environmental Science & Technology, 59(11), 5417–5430. 10.1021/acs.est.4c04557

69. Stockbridge, R. B., & Wackett, L. P. (2024). The link between ancient microbial fluoride resistance mechanisms and bioengineering organofluorine degradation or synthesis. Nature Communications, 15(1), 4593. 10.1038/s41467-024-49018-1

70. Strubel, V., Rast, H. G., Fietz, W., Knackmuss, H.-J., & Engesser, K. H. (1989). Enrichment of dibenzofuran utilizing bacteria with high co-metabolic potential towards dibenzodioxin and other anellated aromatics. FEMS Microbiology Letters, 58(2–3), 233–238. 10.1111/j.1574-6968.1989.tb03050.x

71. Tang, Z., Vogel, T. M., Wang, Q., Wei, C., Ali, M., & Song, X. (2024). Microbial defluorination of TFA, PFOA, and HFPO-DA by a native microbial consortium under anoxic conditions. Journal of Hazardous Materials, 465, 133217. 10.1016/j.jhazmat.2023.133217

72. Thermo Fisher Scientific. (2016, September). *Thermo Scientific Orion Fluoride Ion Selective Electrode—User manual*. https://documents.thermofisher.com/TFS-Assets/LSG/manuals/D15872~.pdf

73. Twigg, L. E., & Socha, L. V. (2001). Defluorination of sodium monofluoroacetate by soil microorganisms from central Australia. Soil Biology and Biochemistry, 33(2), 227–234. 10.1016/S0038-0717(00)00134-6

74. Visscher, P. T., Culbertson, C. W., & Oremland, R. S. (1994). Degradation of trifluoroacetate in oxic and anoxic sediments. Nature, 369(6483), 729–731. 10.1038/369729a0

75. Wackett, L. P. (2021). Why Is the Biodegradation of Polyfluorinated Compounds So Rare? mSphere, 6(5), 10.1128/msphere.00721-21. https://doi.org/10.1128/msphere.00721-21

76. Wackett, L. P. (2022). Nothing lasts forever: Understanding microbial biodegradation of polyfluorinated compounds and perfluorinated alkyl substances. Microbial Biotechnology, 15(3), 773–792. 10.1111/1751-7915.13928

77. Wackett, L. P. (2024). Evolutionary obstacles and not C–F bond strength make PFAS persistent. Microbial Biotechnology, 17(4), e14463. 10.1111/1751-7915.14463

78. Wick, R. (2026). *Rrwick/Porechop* [C++]. https://github.com/rrwick/Porechop (Original work published 2017)

79. Zheng, G., Eick, S. M., & Salamova, A. (2023). Elevated Levels of Ultrashort- and Short-Chain Perfluoroalkyl Acids in US Homes and People. Environmental Science & Technology, 57(42), 15782–15793. 10.1021/acs.est.2c06715

