## Supplementary material for "TFA-defluorinating microbial consortium achieved with analogue enrichment"

#### Calibration curve for quantification of free $F^-$ with the fluoride ion-specific electrode

*Supplementary table S1. Overview of the concentrations used to prepare a calibration curve in the 0.5-25* µM *range.*

| 1 mM $F^-$ volume added (µl) | Total 1 mM $F^-$ volume added (µl) | Resulting µM | Resulting ppm |
|---|---|---|---|
| 10 | 10 | **0.53** | 0.01 |
| 10 | 20 | **1.05** | 0.02 |
| 20 | 40 | **2.11** | 0.04 |
| 20 | 60 | **3.16** | 0.06 |
| 40 | 100 | **5.27** | 0.1 |
| 200 | 300 | **15.27** | 0.29 |
| 200 | 500 | **25.28** | 0.48 |

The electrode exhibits Nernstian behaviour, meaning there is a log-linear relationship between the potential (mV) and the $F^-$ activity. The theoretical Nernst slope for $F^-$ is -59 mV per decade change in $F^-$ concentration at 25°C.

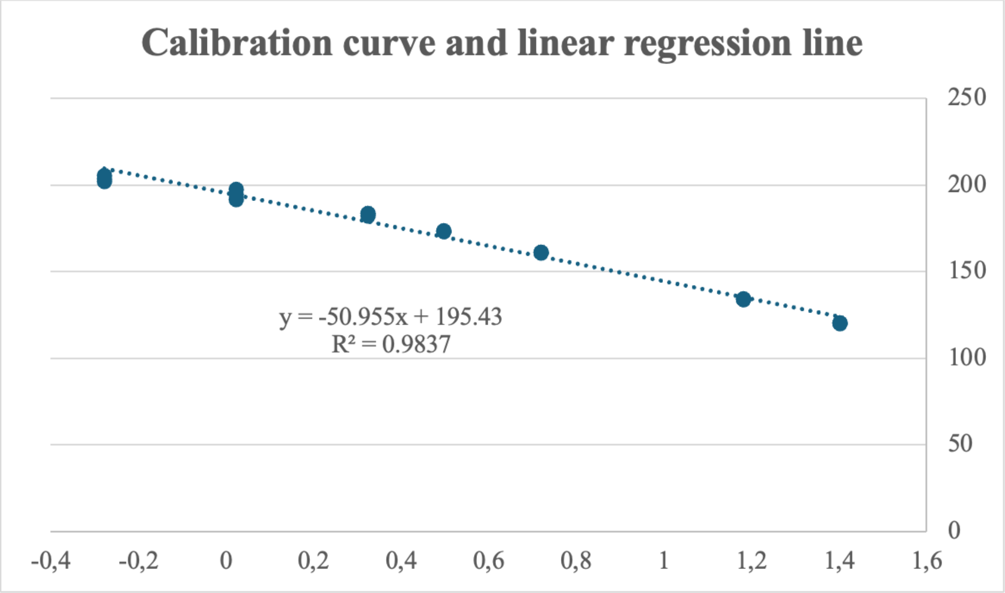


*Supplementary figure S1. Calibration curve prepared according to Thermofisher instrument protocol by standard addition of 1 mM $F^-$ to M9 minimal media mixed 1:1 with low-level TISAB.*

The calibration curve (Figure S1) shows a good linear relationship with an $R^2$ value of 0.9837. The curve also shows a slightly sub-Nernstian electrode response with a slope of 50.995 mV. This indicates a slightly lowered sensitivity of the electrode to $F^-$.

### Tests of two-way ANOVA mixed model assumptions

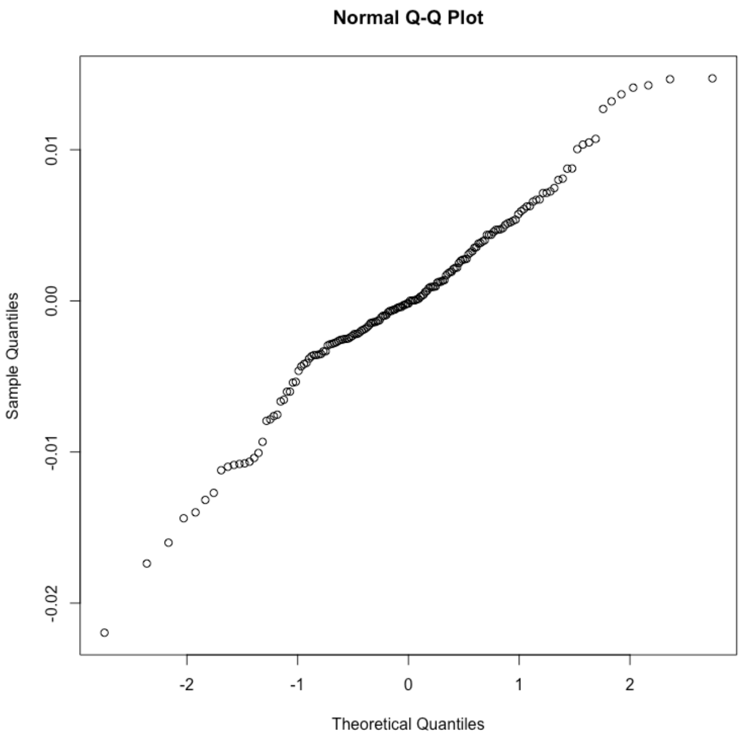


*Supplementary figure S2. QQ-plot of residual values from the two-way ANOVA mixed model*

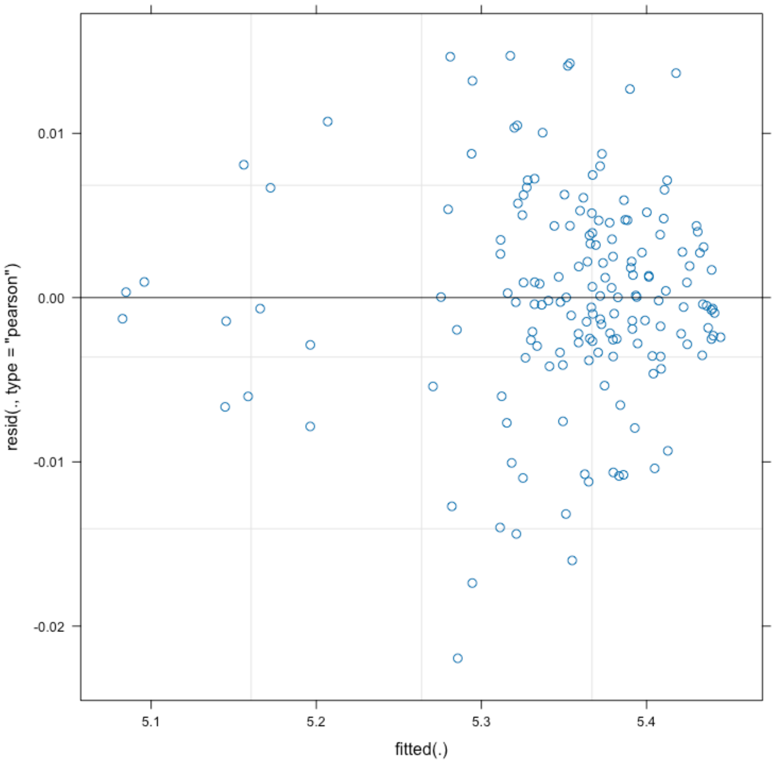


*Supplementary figure S3. Residual plot of residual values plotted against predicted risiduals values from the two-way ANOVA mixed model*

#### Data collected with fluoride ion specific electrode for 30-day experiment

***Supplementary table S2.*** *Samples have been given a shorthand;* ***A****31 is sample DFA1****A*** *on TFA replicate 1, etc.*

| | Days: | 2 | | 5 | | 9 | | 15 | | 30 | |
|---|---|---|---|---|---|---|---|---|---|---|---|
| | | mV | pH | mV | pH | mV | pH | mV | pH | mV | pH |
| Abiotic - MFA | F1 | 222.6 | 5.5 | - | - | 208.3 | 5.5 | 209.8 | 5.5 | 204.3 | 5.5 |
| Abiotic - DFA | F2 | 212.1 | 5.5 | 224.1 | 5.5 | 221.5 | 5.5 | 219.5 | 5.5 | 221.6 | 5.5 |
| Abiotic - TFA | F3 | 212.7 | 5.5 | 220.9 | 5.5 | 217.4 | 5.5 | 209.7 | 5.5 | 205.5 | 5.5 |
| Negative biological | G1 - | 214.1 | 6.5 | 216.5 | 7.5 | 220.2 | 8.5 | 216 | 8.5 | 213.8 | 8.5 |
| Negative biological | G2 - | 208.3 | 6.5 | 219.6 | 8.0 | 224.1 | 8.5 | 222 | 8.5 | 213 | 8.5 |
| Negative biological | G3 - | 212.9 | 6.5 | 219.7 | 8.0 | 225.3 | 8.5 | 219.1 | 8.5 | 213.2 | 8.5 |
| Positive biological (0.01ppm) | G1 + | 215.6 | | 213.1 | | 220.3 | | 212.9 | 7.0 | 210.5 | 7.0 |
| Positive biological (0.02 ppm) | | 212.8 | | 209.4 | | 215.3 | | 210.5 | 7.0 | 207.8 | 7.0 |
| Positive biological (0.04 ppm) | | 210.0 | | 205.8 | | 211.8 | | 206.7 | 7.0 | 201.9 | 7.0 |
| Positive biological (0.06 ppm) | | 207.9 | | 201.6 | | 206.6 | | 203 | 7.0 | 198.1 | 7.0 |
| Positive biological (0.1 ppm) | | 203.0 | | 195.8 | | 199.2 | | 197.2 | 7.0 | 193.7 | 7.0 |
| Positive biological (0.29 ppm) | | 188.7 | | 181.5 | | 182.4 | | 181.4 | 7.0 | 183.8 | 7.0 |
| Positive biological (0.48 ppm) | | 178.5 | | 171.0 | | 169.4 | | 168.2 | 7.0 | 171.0 | 7.0 |
| Positive biological (0.29 ppm) | G2+ | 189.0 | | 179.5 | | 184.1 | | 181.6 | 7.0 | 184.2 | 7.0 |
| Positive biological (0.29 ppm) | G3+ | 189.0 | | 180.4 | | 183.6 | | 182.6 | 7.0 | 183.6 | 7.0 |
| | A31 | 210.9 | 6.5 | 214.7 | 7.0 | 203.2 | 7.0 | 201.7 | 7.0 | 206.4 | 7.0 |
| | A32 | 212.0 | 7.0 | 215.1 | 7.0 | 207.1 | 7.0 | 207.5 | 7.0 | 206.2 | 7.0 |
| | A33 | 214.1 | 6.5 | 216.1 | 7.0 | 210.0 | 7.0 | 208.4 | 7.0 | 206.6 | 7.0 |
| **TFA** | B31 | 216.2 | 6.0 | 216.9 | 7.0 | 214.4 | 7.0 | 208.9 | 7.0 | 207.7 | 7.0 |
| | B32 | 217.5 | 6.0 | 217.5 | 7.0 | 208.1 | 7.0 | 210.2 | 7.0 | 206.8 | 7.0 |
| | B33 | 216.9 | 6.0 | 217.6 | 7.0 | 211.3 | 7.0 | 212.0 | 7.0 | 207.7 | 7.0 |
| | C31 | 215.4 | 6.5 | 206.9 | 7.0 | 184.5 | 7.0 | 174.9 | 7.0 | 163.5 | 7.0 |

| | | | | | | | | | | | |
|---|---|---|---|---|---|---|---|---|---|---|---|
| | C32 | 216.1 | 6.0 | 203.4 | 7.0 | 179.2 | 7.0 | 170.4 | 7.0 | 161.0 | 7.0 |
| | C33 | 217.4 | 5.5 | 199.8 | 7.0 | 180.1 | 7.0 | 171.4 | 7.0 | 161.6 | 7.0 |
| | D31 | 216.9 | 6.0 | 209.4 | 7.0 | 201.9 | 7.0 | 193.2 | 7.0 | 199.5 | 7.0 |
| | D32 | 219.9 | 6.0 | 211.9 | 7.0 | 205.7 | 7.0 | 200.9 | 7.0 | 197.0 | 7.0 |
| | D33 | 219.6 | 6.5 | 214.4 | 7.0 | 207.3 | 7.0 | 201.9 | 7.0 | 194.3 | 7.0 |
| | E31 | 219.9 | 6.0 | 215.3 | 7.0 | 214.0 | 7.0 | 210.1 | 7.0 | 201.6 | 7.0 |
| | E32 | 219.2 | 6.0 | 215.0 | 7.0 | 215.1 | 7.0 | 209.6 | 7.0 | 203.6 | 7.0 |
| | E33 | 220.1 | 6.0 | 216.2 | 7.0 | 214.3 | 7.0 | 212.3 | 7.0 | 206.0 | 7.0 |
| | A21 | 229.2 | 6.0 | 214.7 | 6.5 | 220.0 | 7.0 | 211.4 | 7.0 | 206.8 | 7.0 |
| | A22 | 229.4 | 6.0 | 219.7 | 6.5 | 220.9 | 7.0 | 215.4 | 7.0 | 205.0 | 7.0 |
| | A23 | 229.7 | 6.0 | 219.9 | 6.5 | 222.0 | 7.0 | 217.0 | 7.0 | 205.9 | 7.0 |
| | B21 | 229.0 | 5.5 | 222.2 | 7.0 | 222.9 | 7.0 | 217.6 | 7.0 | 206.9 | 7.0 |
| | B22 | 229.3 | 6.0 | 222.4 | 6.5 | 222.0 | 7.0 | 214.1 | 7.0 | 195.8 | 7.0 |
| | B23 | 230.0 | 5.5 | 228.5 | 6.5 | 224.1 | 7.0 | 216.4 | 7.0 | 203.2 | 7.0 |
| | C21 | 229.3 | 6.5 | 221.3 | 7.0 | 214.4 | 7.0 | 193.5 | 7.0 | 172.9 | 7.0 |
| **DFA** | C22 | 229.5 | 6.0 | 224.8 | 6.5 | 214.7 | 7.0 | 197.4 | 7.0 | 177.5 | 7.0 |
| | C23 | 228.2 | 6.0 | 223 | 6.5 | 216.1 | 7.0 | 195.5 | 7.0 | 175.1 | 7.0 |
| | D21 | 229.9 | 6.0 | 221.4 | 6.5 | 218.1 | 7.0 | 211.0 | 7.0 | 206.5 | 7.0 |
| | D22 | 230.3 | 6.5 | 222.5 | 6.5 | 221.4 | 7.0 | 214.7 | 7.0 | 204.5 | 7.0 |
| | D23 | 231.0 | 6.0 | 225.8 | 6.5 | 222.6 | 7.0 | 215.9 | 7.0 | 202.0 | 7.0 |
| | E21 | 230.7 | 6.0 | 227.1 | 6.5 | 225.6 | 7.0 | 213.4 | 7.0 | 208.6 | 7.0 |
| | E22 | 230.1 | 6.0 | 226.3 | 6.5 | 226.9 | 7.0 | 213.7 | 7.0 | 210.3 | 7.0 |
| | E23 | 230.5 | 5.5 | 227.7 | 7.0 | 226.3 | 7.0 | 215.3 | 7.0 | 207.9 | 7.0 |
| | A11 | 196.0 | 5.5 | 183.9 | 6.0 | 165.4 | 6.5 | 131.6 | 7.0 | 95.9 | 7.0 |
| **MFA** | A12 | 201.5 | 6.0 | 186.9 | 6.0 | 164.0 | 6.5 | 108.5 | 7.0 | 83.9 | 7.0 |
| | A13 | 203.7 | 6.0 | 189.9 | 6.0 | 171.2 | 6.5 | 123 | 7.0 | 90.7 | 7.0 |
| | B11 | 200.2 | 6.0 | 175 | 6.5 | 147.2 | 6.5 | 102.6 | 7.0 | 85.6 | 7.0 |

| | | | | | | | | | | |
|---|---|---|---|---|---|---|---|---|---|---|
| B12 | 204.1 | 6.0 | 191.2 | 6.0 | 173.9 | 6.5 | 127.6 | 7.0 | 79.3 | 7.0 |
| B13 | 206 | 6.0 | 185.1 | 6.0 | 165.1 | 6.5 | 118.1 | 7.0 | 82.2 | 7.0 |
| C11 | 208.4 | 6.0 | 190.5 | 6.0 | 165.5 | 6.5 | 118.3 | 7.0 | 85 | 7.0 |
| C12 | 209.4 | 6.0 | 189.3 | 6.0 | 165.3 | 6.5 | 128.5 | 7.0 | 92.5 | 7.0 |
| C13 | 207.4 | 6.0 | 186.2 | 6.0 | 165.5 | 6.5 | 136.2 | 7.0 | 92.7 | 7.0 |
| D11 | 211.1 | 5.5 | 192.8 | 6.0 | 172.2 | 6.5 | 140.1 | 7.0 | 96.3 | 7.0 |
| D12 | 214.3 | 5.0 | 201 | 5.5 | 183.1 | 6.5 | 172.3 | 7.0 | 124.1 | 8.5 |
| D13 | 201.7 | 6.0 | 170.9 | 6.5 | 141.1 | 7 | 94.7 | 7.0 | 75.5 | 7.0 |
| E11 | 210 | 6.0 | 187.8 | 6.0 | 167.4 | 6.5 | 135.6 | 7.0 | 84.8 | 7.0 |
| E12 | 214 | 6.0 | 192 | 6.0 | 168.8 | 6.5 | 132.8 | 7.0 | 83.4 | 7.0 |
| E13 | 212.8 | 5.5 | 192.1 | 6.0 | 169.8 | 6.5 | 144.4 | 7.0 | 87.8 | 7.0 |
| **Negative control for MFA** | 214.3 | | 217.5 | | 212.4 | 7.5 | 201.9 | 8.0 | 203.2 | 8.0 |

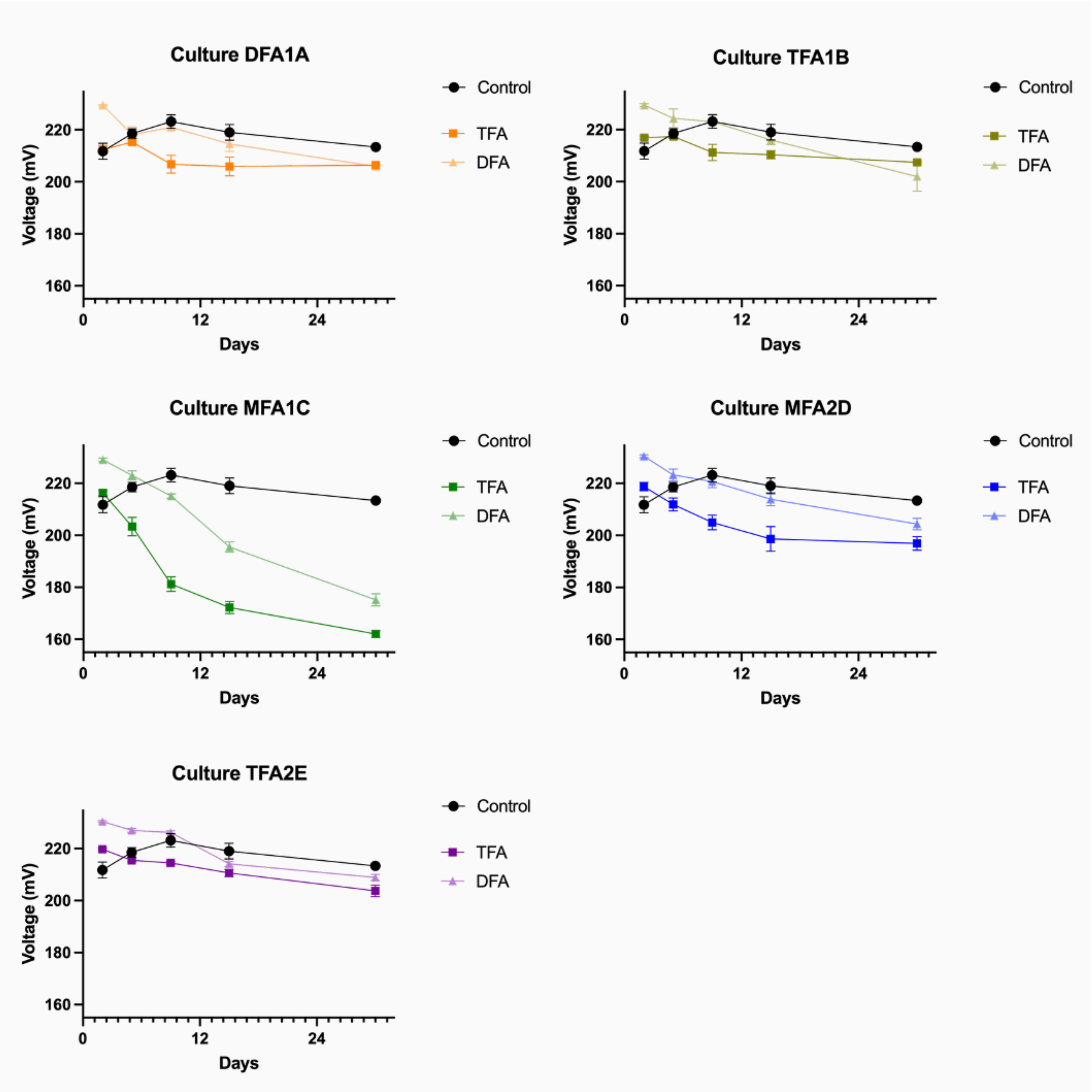


*Supplementary figure S4. Average voltage measurements of triplicates incubated with DFA and TFA across the 30-day experiment. A lower voltage reading is associated with greater $F^-$ release.*

*Supplementary table S3. Calculation of total fluoride release from culture MFA1C.*

| | |
|---|---|
| Converting mV to μM using the calibration curve shown in Fig. S1 | C31: $10^{\frac{163.5\ mV-195.43}{-50.85}} = 4.23\ \mu M\ F^{-}$<br>C32: $10^{\frac{161.0\ mV-195.43}{-50.85}} = 4.74\ \mu M\ F^{-}$<br>C33: $10^{\frac{161.6\ mV-195.43}{-50.85}} = 4.61 \mu M\ F^{-}$ |
| Correcting for dilution factor of 2.75 (400 μL in a final volume of 1100 μL) | C31: $4.23\ \mu M\ F^{-} \cdot 2.75 = 11.64\ \mu M\ F^{-}$<br>C32: $4.74\ \mu M\ F^{-} \cdot 2.75 = 13.03\ \mu M\ F^{-}$<br>C33: $4.61\ \mu M\ F^{-} \cdot 2.75 = 12.60\ \mu M\ F^{-}$ |
| Substracting background from negative control sample | Average negative control: $1.22\ \mu M\ F^{-}$<br>C31: $4.23\ \mu M\ F^{-} \cdot 2.75 = 10.42\ \mu M\ F^{-}$<br>C32: $4.74\ \mu M\ F^{-} \cdot 2.75 = 11.81\ \mu M\ F^{-}$<br>C33: $4.61\ \mu M\ F^{-} \cdot 2.75 = 11.46\ \mu M\ F^{-}$ |
| Average fluoride release from sample MFA1C<br>Standard deviation | |
| Percentage of theroetical maximum fluoride release from 33 mM TFA | |

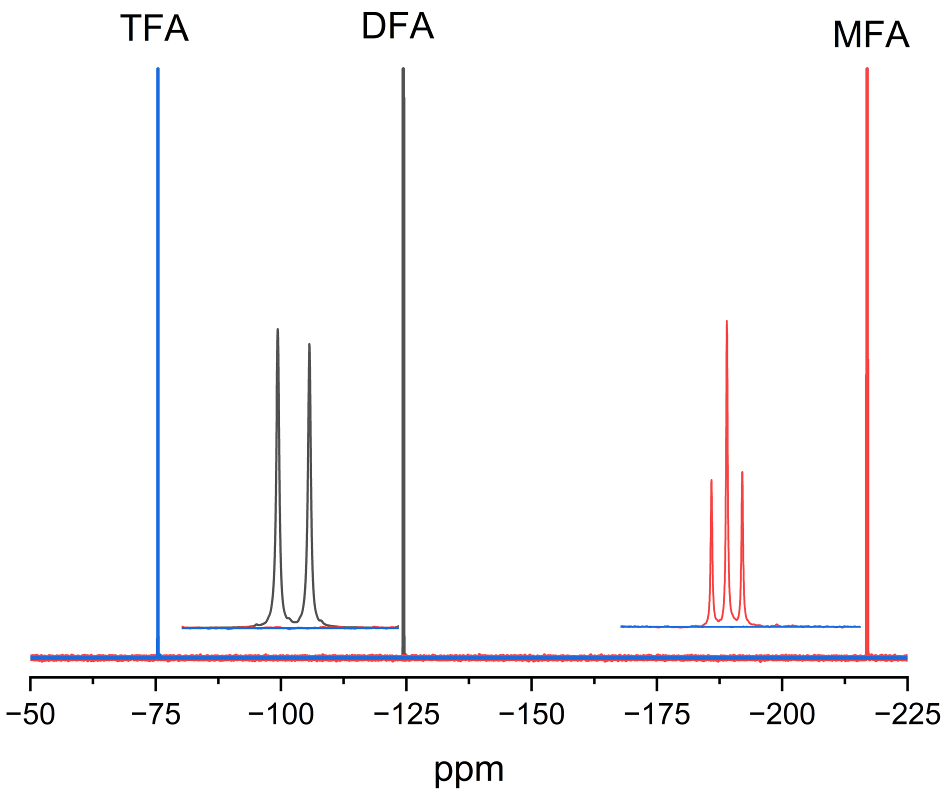


***Supplementary figure S5.*** *$^{19}$F-NMR reference spectra of TFA (blue), DFA (black), and MFA (red), acquired under the same conditions as the culture supernatants. Insets show expansions of the DFA doublet and the MFA triplet, the multiplicities of which arise from coupling to the adjacent proton(s) and which were used to assign the corresponding signals in culture MFA1C.*
